# The motor apparatus of head movements in the Oleander hawkmoth (*Daphnis nerii*, Lepidoptera)

**DOI:** 10.1101/2023.08.15.553342

**Authors:** Agnish D. Prusty, Sanjay P. Sane

## Abstract

Head movements of insects play a vital role in diverse locomotory behaviors including flying and walking. Because insect eyes move minimally within their sockets, their head movements are essential to reduce visual blur and maintain a stable gaze. As in most vertebrates, gaze stabilization behavior in insects requires integration of both visual and mechanosensory feedback by the neck motor neurons. Whereas visual feedback is derived from the optic flow over the retina of their compound eyes, mechanosensory feedback is derived from their organs of balance, similar to the vestibular system in vertebrates. In Diptera, vestibular feedback is derived from the halteres – modified hindwings that evolved into mechanosensory organs – and is integrated with visual feedback to actuate compensatory head movements. However, non-Dipteran insects including Lepidoptera lack halteres. In these insects, vestibular feedback is obtained from the antennal Johnston’s organs but it is not well-understood how it integrates with visual feedback during head movements. Indeed, although head movements are well-studied in flies, the underlying motor apparatus in non-Dipteran taxa has received relatively less attention. As a first step towards understanding compensatory head movements in the Oleander hawkmoth *Daphnis nerii*, we image the anatomy and architecture of their neck joint sclerites and muscles in using X-ray microtomography, and the associated motor neurons using fluorescent dye fills and confocal microscopy. Based on these morphological data, we propose testable hypotheses about the putative function of specific neck muscles during head movements, which can shed light on their role in neck movements and gaze stabilization.

## 1. Introduction

Most vertebrate animals, including humans, track moving objects by moving their eyes or adjusting their head (Land, 1999). In insects however, the eye movements relative to the head are very limited (Burtt & Patterson, 1970; Fenk *et. al*., 2022), and hence gaze stabilization is achieved largely through head movements, thereby reducing motion blur and keeping the salient features of the environment steady on their retina (Hardcastle & Krapp, 2016). Because head movements play a crucial role in a vast range of behaviors in insects, they have been studied in diverse insects including blowflies (Land, 1999; Kress & Egelhaaf, 2012; Hengstenberg, 1993), praying mantids (Liske & Mohren, 1984; Rossel, 1980), moths (Chatterjee et al. 2022, Dombrowski et al., 1990), bees (Boeddeker & Hemmi, 2010), ants (Raderschall, Narendra & Zeil, 2016), and wasps (Stürzl, *et.al.* 2016). These behaviors are wide-ranging and encompass flight, foraging, pursuing prey, evading predators, constructing or safeguarding nests, as well as reproductive activities such as mate selection, courtship, and oviposition. These diverse insect species provide a rich contextual foundation for studying the significance of head movements.

Head movements in insects are actuated by the neck-thorax or cervical joint that houses the cervical connective nerve, which transmits neural information between the brain and the thoracic ganglia. Spanning this joint are several muscles which actuate head movements around four-axes, including three rotational (yaw, roll, and pitch) axes and one translational axis allowing the neck to extend or withdraw (Snodgrass, 2018). These muscles are activated by motor neurons residing in the head and the prothoracic ganglia, requiring fine coordination of their activity across segments. The activity in the motor neurons, in turn, is elicited by feedback from multiple sensory modalities involved in involuntary and voluntary behaviors. Thus, to understand the neuromuscular basis of head movements, it is essential to characterize the apparatus that actuates the neck joint in diverse insects. This includes the description of their muscles, the related sclerites, the underlying motor neurons in addition to the joint motion. The insect neck musculo-skeletal and neuro-muscular apparatus has been described at length in blowflies (Diptera) (Sandeman & Markl, 1979; Strausfeld *et al*, 1987; Milde et al, 1987; Gilbert et al, 1995), honeybees (Hymenoptera) (Goodman *et al*, 1987; Schröter *et al*, 2007; Berry & Ibbotson, 2010; Hung et al 2011), crickets and locusts (Orthoptera) (Shepheard, 1973, 1974; Kien, 1980; Honneger, Altman *et al*, 1984; Bartos & Honneger, 1992; Baader 1991) but to a lesser extent in moths (Lepidoptera) (Eaton 1971; Bharadwaj *et al*, 1974).

Recent work has demonstrated that compensatory head movements in hawkmoths are essential for stable flight, and also that the neck motor system integrates multimodal information to generate these movements (Chatterjee et al, 2022). A detailed investigation of their motor apparatus is, however, prerequisite towards gaining a mechanistic understanding of this multisensory integration. Recent tools such as X-ray Microtomography (Micro-CT) and fluorescent dye fill techniques combined with confocal microscopy afford us vastly better resolution of the neck motor apparatus than has been previously possible. Here, we use these techniques to outline the 3D architecture and morphology of the neck motor apparatus in the Oleander hawk moth (*Daphnis nerii*) including the neck skeletal elements and the muscles that actuate them. We additionally describe the neuroanatomy of the neck motor neurons of these muscles through fluorescent dye-fills. Together, these data will help in future studies that seek to understand the multisensory integration of head movements for gaze stabilization and other behaviors in hawkmoths and other insects.

## 2. Materials and Methods

### 2.1 Moth culture

The Oleander hawkmoth *Daphnis nerii* occurs in Bangalore and also widely across the peninsular India. The Oleander hawkmoths used in this study were bred in their natural day-night cycle and housed in the greenhouse facility at National Centre for Biological Sciences, Bangalore, India. The ambient conditions with average temperature of around 25 – 28^°^C and a relative humidity of about 70 – 85 % were maintained in the breeding and culture rooms. The moth larvae were raised on a diet of fresh, tender leaves from the *Nerium oleander* plant in the 1^st^ and 2^nd^ instar phases, and on mature leaves in 3^rd^ or higher instars. The larvae were kept in plastic mesh-topped boxes for air ventilation and light passage. After the feeding behaviour stopped, the 5^th^ (and last) instar larvae were moved into a plastic box with vermiculite and saw-dust bedding, under which they burrowed before pupation. A subset of the one-day old males and females post were released in the breeding cage whereas the remaining were used for experiments. The eggs laid by the mated females on the *Nerium* leaves were collected for further culturing.

### 2.2 3D-reconstruction of neck arthrology using microtomography

The moth was first cold-anesthetized and its abdomen, wings and palps were removed. The antennae were next cut to reduce their extraneous length and the head, neck and thorax were descaled. After removing the ventral part of the episternum, a small window was cut in the head cuticle to allow better penetration of the dye and staining of the neck muscles. We fixed the tissues in 4% paraformaldehyde (PFA) solution in phosphate buffer solution (PBS), at room temperature (25°C) in dark for about 12-14 hours. The sample was then dehydrated through an ascending series of ethanol solution (10 to 90% in steps of 10%, and twice in 100%) for 10 minutes each. The sample was next immersed with 0.25% of phosphotungstic acid (PTA) in ethanol solution and placed on a gentle shaker for staining at room temperature over five days. After this, the sample was dried using carbon dioxide critical point drying (Leica EM CPD 300 Critical Point Dryer, Leica microsystems, Wetzlar, Germany) at a slow speed setting and CO2 gas was exchanged out at slow heating for 18 cycles. The sample was then mounted on a holder and imaged using the X-ray Microtomography (Micro-CT) scanner (SkyScan 1272, Bruker, Belgium) at 40 kV, power of 10 W, exposure time of 699 ms and a resolution of 2 μm. The X-ray computed tomography images were then reconstructed using the SkyScan volumetric NRecon software. We traced and interpolated the neck muscles and relevant portions of the moth skeleton voxel data using the Amira 6.2 software (FEI Visualization Sciences Group, France and Zuse Institute, Germany) to segment out the three-dimensional model of the neck motor apparatus. The interactive pdf (Supplementary Figure 1) of the 3-D model was made using Adobe Pro (Adobe Systems; Mountain View, CA, USA).

### 2.3. Neuroanatomy

#### Preparation of sample

The moths were anaesthetized at -23°C for about 10 minutes and inserted in a cylindrical tube. Their thorax was fixed to the tube wall using dental wax. Their neck region was then descaled with a fine brush to prepare the moths for surgery. After removal of the ventral episternal cuticle, we could access two ventral neck muscles (cv1, v) and one tentorial muscle t2 but not the ventral muscle cv2 and tentorial muscle t1 (see Results) without loss of haemolymph due to bleeding. To access the dorsal neck muscles, a small window was cut out on the dorsal cervix by removing a tiny portion of the soft tissue flap which connects the post-occipital ridge in posterior head-capsule to the pronotum in the anterior part of the prothorax. As this dissection was more prone to bleeding due to several haemolymph vessels underlying this region, only three dorsal neck muscles (the dorsal muscles d1-3; see Results) could be accessed during this surgery.

#### Motor neuronal labelling

Next, we selectively punctured the neck muscles using a fine syringe needle or a minuten pin to damage the motor neuronal termini within the muscles, and inserted the water-soluble fluorophore dyes (Invitrogen, Thermo-Fisher Scientific, Massachusetts, USA) into these muscles. The dye was taken up and actively transported across the neuron. We specifically used dyes including Texas Red Dextran (D3328, zwitterionic, lysine fixable, 3000 molecular weight (MW), excitation at 595 nm & emission at 615 nm) and Alexa Fluor Dextran 488 (D22910, anionic, fixable, 10000 MW, excitation at 495 nm & emission at 519 nm) that labeled the cytosol of the neck motor neurons. To allow for a complete retrograde labeling of these motor neurons, we kept the moths alive for 48 hours by feeding them 25% sugar solution. The moth was then fully descaled, euthanized, and its extraneous appendages were removed. The tissues were fixed in 4% PFA solution in Phosphate Buffer Solution (PBS), at 4°C for 24 hours or at room temperature (25°C) for about 12-14 hours.

#### Dissection and imaging

We next dissected the brain and the thoracic ganglia of the tissue fixed moth in insect saline (Supplementary materials). The brain and thoracic ganglia samples were washed in phosphate-buffered saline (PBS) and dehydrated by passing them through a sequential series of ethanol dilutions (50%, 60%, 75%, 90% and twice in 100%; 10 minutes in each concentration). To clear the samples, we placed the dehydrated tissues in methyl salicylate and mounted them on glass cover slips. The stained tissues were then scanned and imaged with appropriate laser excitations (488 nm He–Ne laser or 543 nm Kr-Ar laser) under 10× and 20× objectives using a laser-scanning confocal microscope (Olympus FV1000, FV 3000, Microscopy Technologies, Evident Corporation, Shinjuku-ku, Tokyo, Japan). To visualize and stack the scanned raw images, we used Olympus Fluo-view software (FV-10 ASW Version 4.2b) and processed for background subtraction or signal contrast enhancement using the Fiji ImageJ software (Version 1.53c, Wayne Rasband, NIH, USA).

#### 3D rendering of the neck skeletal and muscle structures

The detailed rendering of the 3D structure of the neck skeleton and muscles using X-ray microtomography provided a clear picture of the joints, their articulation in the neck skeleton, the geometry and orientation of the muscles, and attachment points in the cuticular segments. Together, this information offers insights into how the muscle contractions generate the forces that actuate the various head movements. It also allowed us to generate a 3-D interactive view for the head capsule, cuticular arrangements in head-neck-prothoracic segments and neck muscles (Supplementary Figure 1). The micro-CT imaging was complemented by microdissections of the neck region to remove the muscles and soft tissues to observe the skeletal parts under the microscope. The posterior cuticle of the head capsule, the neck and the anterior part of the prothoracic exoskeleton (with invaginations) are especially crucial for the understanding of the neck motor apparatus. In this paper, we have largely retained the terminology and conventions for anatomical nomenclature for head and thorax described for the tobacco hornworm hawkmoth *Manduca sexta* (Eaton, 1971).

## 3. Results

### 3.1 Neck arthrology

The cervical (neck) joint is comprised of the posterior cuticle of the head capsule, the neck and the anterior part of the prothoracic exoskeleton, with invaginations (Figure 1a). The outer covering of the neck is soft and membranous, thereby enabling easy movement of the head relative to the thorax. The dorsal part of this membranous region contains bilaterally symmetric, soft, flap-like structures which lie between the pronotum and the head. Large paired trachea and the oesophagus are located dorsal to the cervical nerve. The paired trachea passes through the dorsal occipital foramen in the head capsule, whereas the gut and the nerve cord project through the foramen magnum (Figure 1b).

**Figure 1.**
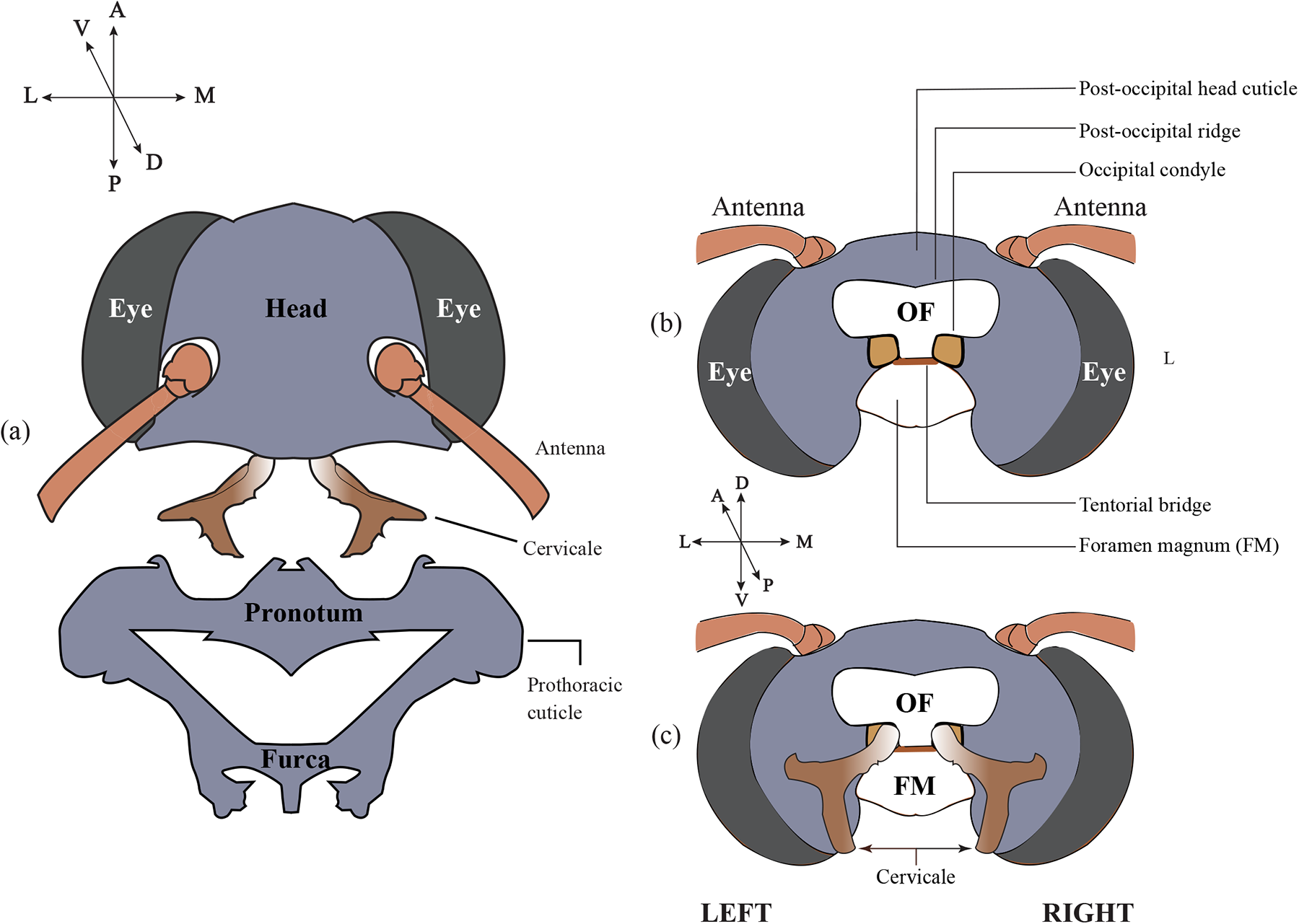
Neck arthrology. A cartoon describing the skeletal structures and joint articulations in the head, neck and prothoracic segments. Grey color marks the cuticle, and graded brown marks the cervicale. Here and in all subsequent figures, the frame of reference is presented adjacent to the figure. A=anterior, P=posterior, D=dorsal, V=ventral, M=medial and L=lateral. (a) Dorsal view of the head, the cervical sclerite (cervicale; graded brown) and the anterior part of prothorax. (b) Posterior view of the head indicating the relevant locations for neck musculoskeletal architecture such as the occipital structures including the occipital condyle (yellow) into which one end of the cervicale inserts to form the pivot. The tentorial bridge (red) and the foramen magnum (FM, unfilled) are also depicted. (c) Posterior view of the head with the cervicale (graded brown) articulating at the occipital condyles (yellow) as sockets. The cuticle of head, pronotum and Furca 1 is shown in grey.

#### Cervicale

A cervical sclerite, henceforth called the *cervicale* (Eaton, 1971), lies within the neck and supports the head capsule with respect to the thorax. It forms the intermediate attachment point within the neck for the neck muscles from the prothorax as well as the head (Figure 1a, c). The cervicale is a bilaterally symmetric structure with its pair of anterior tips forming the anchor points on which the head capsule rests. The socket-like grooves at the occipital condyles in the posterior head-capsule form the pivots, about which the entire head rotates depending on the action of neck muscles (Figure 1b, Supplementary Figure 2a). The structure of cervicale appears to have graded sclerotization, as evident from the changes in chitin-based brown color profile (Supplementary Figure 2). At the anterior end, the cervicale is lighter suggesting a softer, less sclerotized tissue. Perhaps due to reduced sclerotization, this tissue is tough but partially elastic and flexible, similar to tendons. At the posterior end where it contacts the prothorax, the cervicale cuticle is deep brown and more sclerotized, and mechanically hard and rigid. This posterior-ventral part of the cervicale is connected to the prothoracic ventral episternum through a soft tissue flap.

### 3.2 Neck muscles

Based on their location with respect to the nerve cord, the neck muscle groups are broadly divided into dorsal and ventral. If they attach to the tentorium, they are termed as tentorial. The nomenclature used in this study has been derived from anatomical description in the hawkmoth *Manduca sexta* (Eaton, 1971, 1974) and honeybees *Apis mellifera* (Berry & Ibbotson, 2010).

#### Dorsal neck muscles

The muscles located above the cervical connective part of nerve cord are classified as dorsal neck muscles. These attach to the posterior head capsule *via* the post-occipital ridge. There are six pairs of dorsal neck muscles (d1-6), of which four (d1-3, d6) are dorso-longitudinal in nature whereas the other two (d4, d5) run along the dorso-ventral axis. Three muscle pairs – d1 (blue; Figure 2a-c), d2 (red; Figure 2a-c), d3 (dark yellow; Figure 2a-c) – are located most dorsally in the moth. Each attach laterally at the post-occipital ridge in the back of the head and medially at the prescutum cuticle below the pronotum in prothoracic exoskeleton. Two dorso-ventral pairs d4 and d5 muscles anchor on the anterior arm of the cervicale. The larger of the two, twisted-ribbon shaped muscle d4 (beige yellow; Figure 2d-f) attaches at the medial part, whereas the smaller d5 muscle (teal; Figure 2d-f) joins the head at the lateral part of the post-occipital ridge. The ventral-most dorsal muscle is the long d6 muscle (brown: Figure 2g-i) which, like the d1-3 muscles, is also longitudinal. The posterior end of d6 anchors at the Furca I (or first furca; Eaton, 1971) in the prothoracic skeleton, which is a skeletal invagination in the ventral part of the prothorax above coxal segment of the prothoracic legs (Figure 2g). The anterior tip of d6 connects to the head cuticle at the lateral end of the post-occipital ridge.

**Figure 2.**
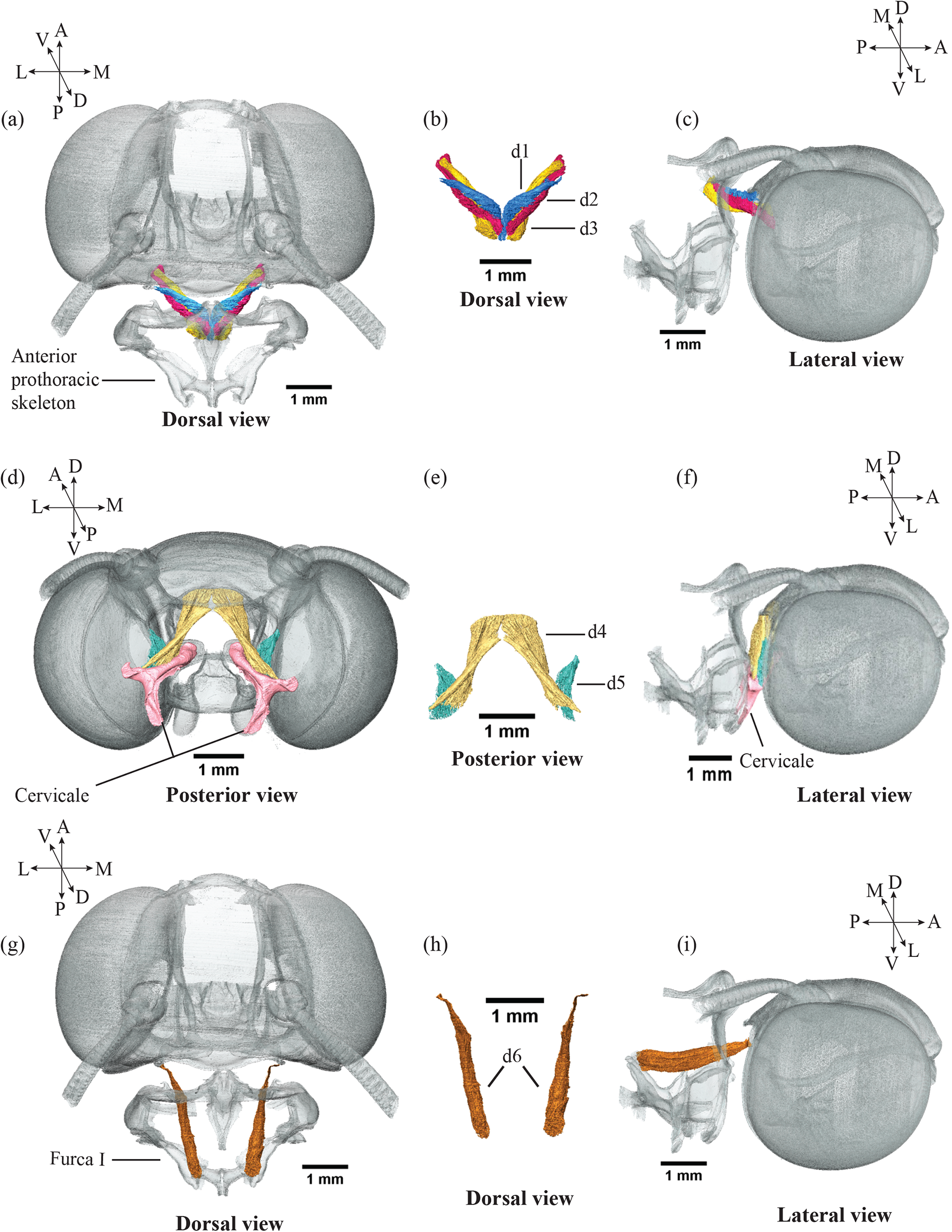
Dorsal neck muscles. (a) Dorsal view of the head and prothoracic skeleton with the three dorsal neck muscles d1 (blue), d2 (red), d3 (dark yellow), and their posterior attachment at the anterior prothorax. (b) A stand-alone dorsal view of the muscles d1-3. (c) lateral view of the three dorsal muscles attaching between the post-occipital ridge of the head and the anterior prothorax. (d) Posterior, (e) stand-alone, and (f) lateral view of the muscle group d4-5 displaying the attachments of the muscles d4 (beige yellow) and d5 (teal) between the post-occipital ridge in the head and the anterior part of the cervicale (pink). (g) Dorsal, (h) stand-alone, and (i) lateral view of muscle d6 (brown) with its attachment at the post-occipital ridge in the head and the Furca I in the thorax. All scale bars are 1 mm

#### Ventral neck muscles

Three pairs of ventral muscles, denoted by v (green; Figure 3a-c), cv1 (violet; Figure 3d-f), and cv2 (orange; Figure 3d-f) are located below the cervical connective of the ventral nerve cord. Whereas the muscle v connects to the head, the other two (cv1-2) attach to the cervicale and are hence only indirectly connected to the head. The ventral muscle v is among the thickest of all the neck muscles (Figure 3a). It is conical in shape, with its broad end attaching at the Furca I in the prothorax. It tapers down sharply with its anterior tip attaching at the ventrolateral end of the foramen magnum. (Figure 3a, c). The other two ventral neck muscles, the cervical ventral muscles (cv1 and cv2) span the Furca I and the cervicale. The cv1 muscle anchors at the ventrolateral end of Furca I and pulls the ventral posterior arm of the cervicale (Figure 3d, f). The cv2 muscle, on the other hand, anchors at the dorsal part of the Furca I and pulls the dorsal-posterior part of the cervical sclerite (Figure 3d, f). The cv1 muscle is almost uniformly broad and thick throughout its length. The cv2 muscle is thick at its prothoracic end but tapers and continues with a thin cross-section to the posterior cervicale attachment site. Anatomically, the ventral muscles v and cv1 are most readily accessible once the ventral episternum cuticle (Supplementary Figure 3) is dissected out.

**Figure 3.**
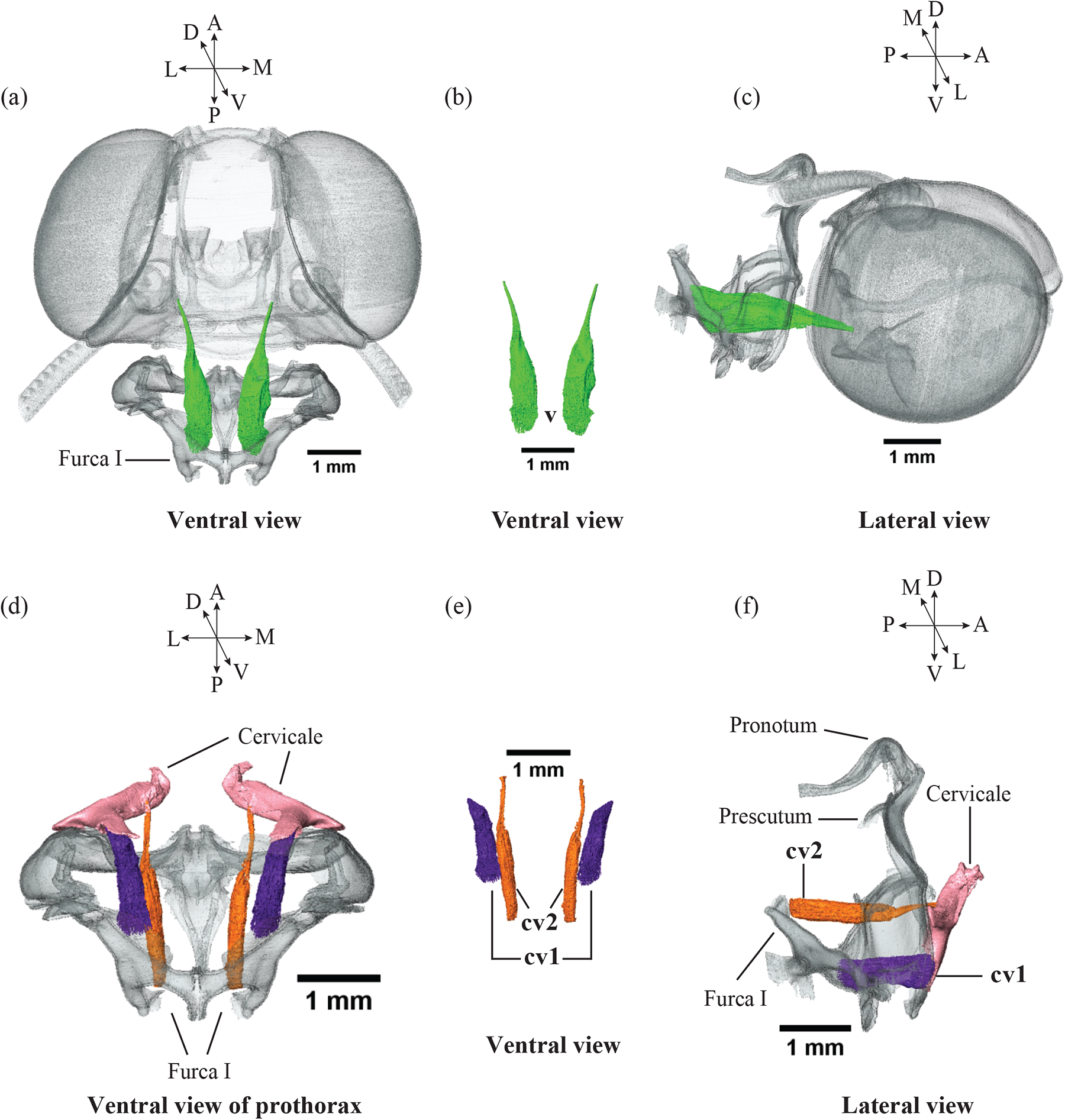
Ventral neck muscles. (a) Ventral, (b) stand-alone, and (c) lateral views of the ventral muscle v (green) located between the Furca I in prothorax and the ventro-lateral position of foramen magnum, below the occipital condyles in the head. (d) Ventral (e) stand-alone and (f) lateral views of the cervical ventral muscle group, cv1 (violet) and cv2 (orange) which attach at the posterior end of the cervicale and at the Furca I in the prothorax. All scale bars are 1 mm

#### Tentorial neck muscles

Muscles that connect the tentorial bridge in the posterior part of the head are called *tentorial neck muscles* (Figure 4). There are two pairs of tentorial muscles: one extending dorsally, and the other ventrally into the thorax to attach onto the prothoracic skeleton (Figure 4a-c). The dorsally extended muscle (t1; bright yellow, Figure 4a-c) is termed as the dorsal tentorial neck muscle and the ventrally extended muscle (t2; pink, Figure 4a-c) as the ventral tentorial neck muscle. The t1 muscle is anchored at the dorsal edge of the anterior episternum whereas the t2 muscle is attached at a ventro-lateral location below the Furca I (Figure 4c). Both t1 and t2 taper and become thinner near the extreme ends whereas they are thicker across the middle of their length. The t1 and t2 muscles anchor at lateral ends in the prothoracic cuticle but connect medially, nearly at the center of the tentorial bridge in the back of the head (Figure 4a).

**Figure 4.**
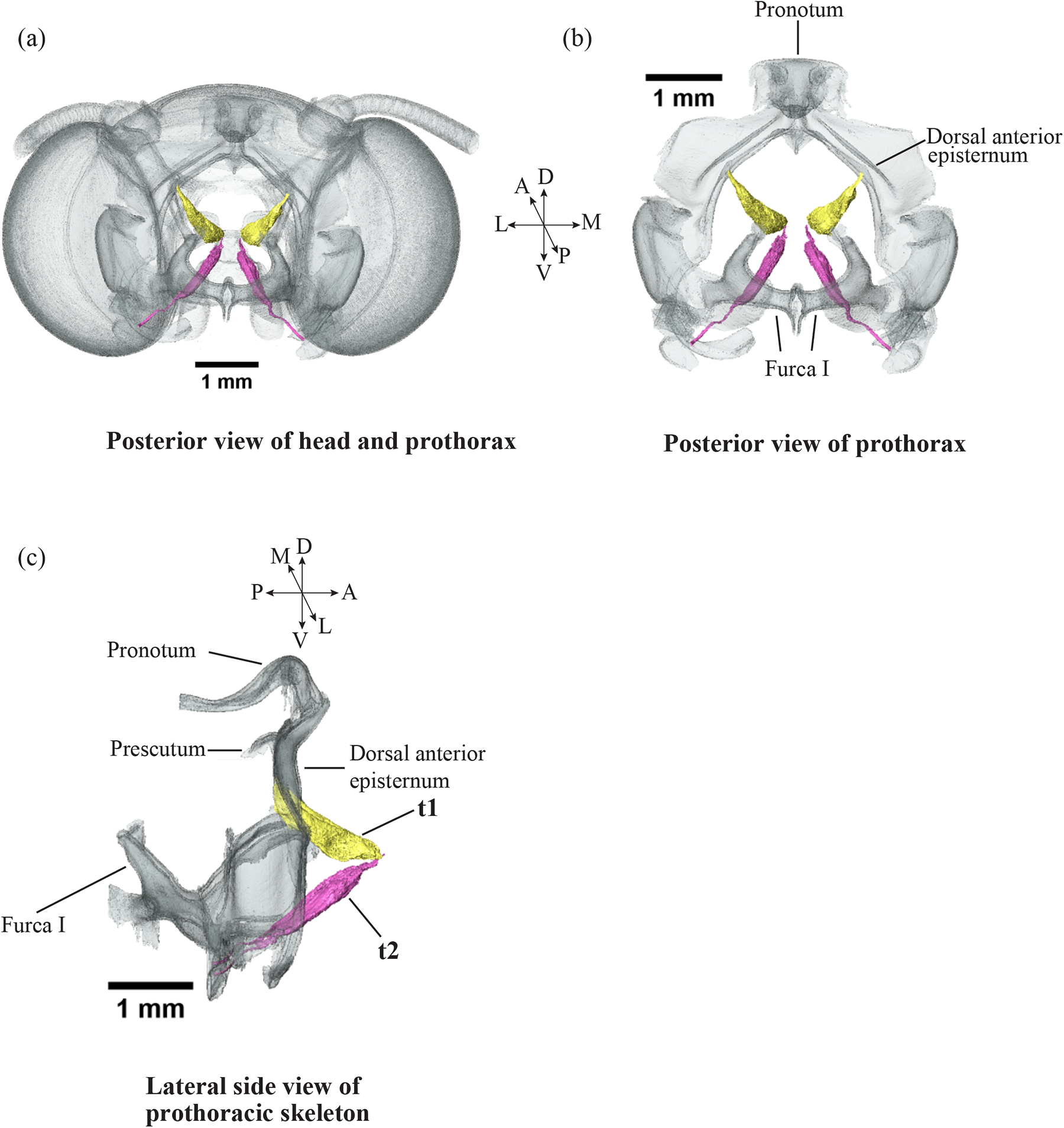
Tentorial neck muscles. (a) Posterior view of the head and prothorax showing the attachment of tentorial neck muscles t1 (bright yellow) and t2 (pink) at the tentorial bridge in the head (b) Posterior view of anchorage points of the neck muscles in the episternum of the prothoracic cuticle. (c) Various locations in the prothorax where the dorsal tentorial muscle t1 attaches at the dorsal anterior edge of episternum and the ventral tentorial muscle t2 at the Furca I. All scale bars are 1 mm

In summary, eleven bilaterally symmetric pairs of neck muscles span the head-cervicale-prothorax joint. Of these, anterior end of nine neck muscles (d1-6, v, t1-2) attach at posterior locations of the head capsule, such as the post-occipital ridge and the tentorial bridge. The other two muscles (cv1-2) connect their anterior tips to the cervicale. The posterior end of two neck muscles (d4-5) are anchored at the cervicale, whereas the other nine muscles (d1-6, v, t1-2) connect to the anterior skeletal segments in the prothorax, and run along the longitudinal anterior-posterior axis starting from head or cervicale towards the prothorax, with variations in the pitch and lateral axes. The neck muscle d4 is anchored laterally on the cervicale but connected medially in the dorsal part of posterior head-capsule, whereas d5 remains lateral throughout the dorso-ventral length. This intricate configuration of the neck muscles and their combinatorial effect enable head movements along all the axes.

### 3.3 Neck motor neurons

#### Motor neurons of the dorsal neck muscles

To identify the motor neurons innervating the neck muscles, we backfilled the muscles with fluorescent dyes (Texas Red; Invitrogen, Thermo-Fisher Scientific, Massachusetts, USA) and imaged the cleared tissues using confocal microscopy. To access the motor neurons innervating the dorsal neck muscles d1-3, we removed the dorsal tissue flap in the neck in both left and right halves. This procedure enabled access to about five to six motor neuronal soma located in the medial region of sub-esophageal zone (SEZ), posterior to the foramen (Figure 5a, b). The axons of these motor neurons project through the sub-esophageal ganglion (SEG) nerve into the dorsal neck muscles (Figure 5c, d). In some fills, the primary neurite projected laterally outward from the medial cell body in the SEZ with its dendrites branching extensively in the antennal mechanosensory and motor center (AMMC) region (Figure 5c, d). Three motor neuronal soma were also observed in the anterior portion of the prothoracic ganglion whose axons descended through anterior prothoracic ganglionic nerves (Figure 5e).

**Figure 5.**
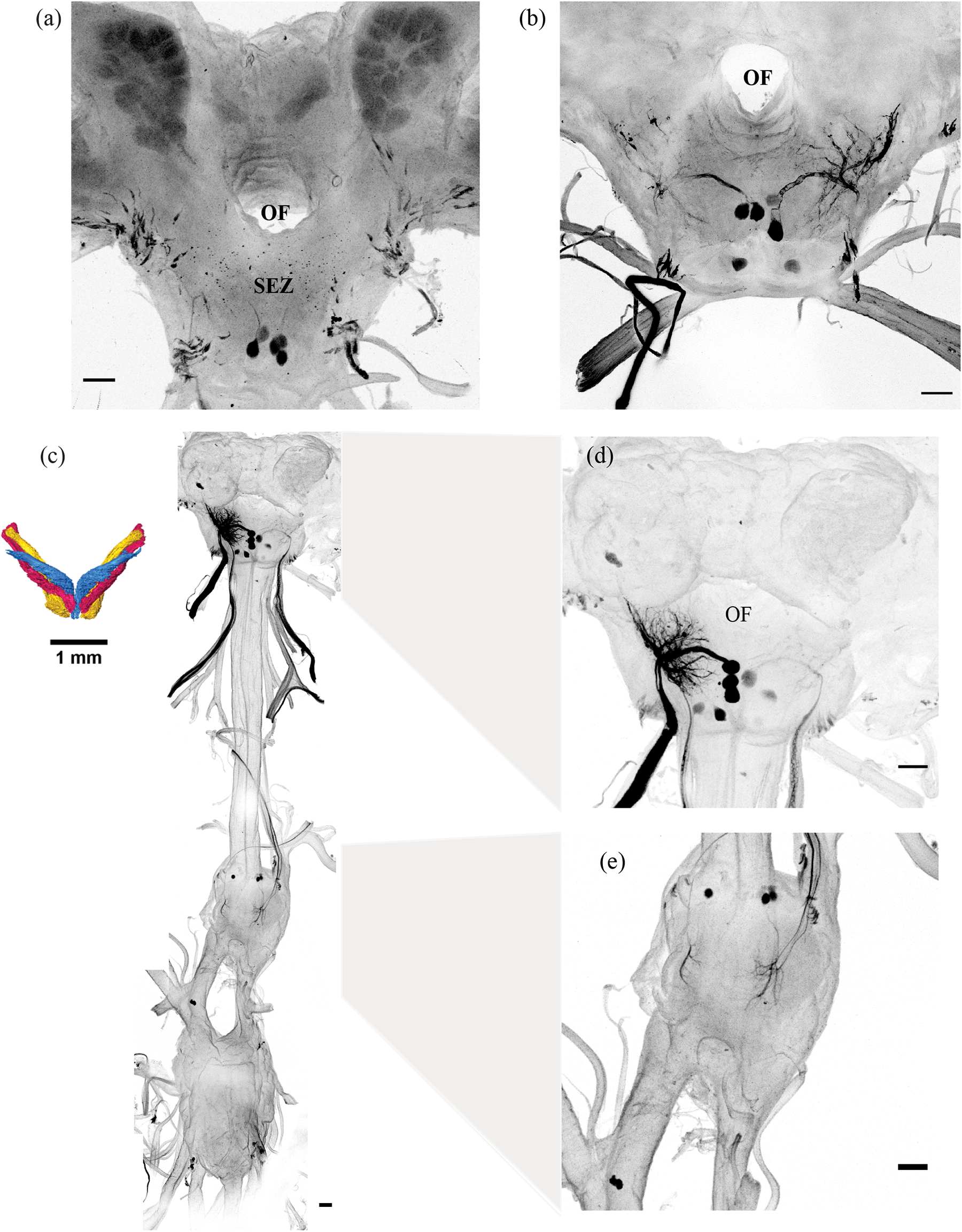
Motor neurons innervating the dorsal neck muscles. (a) Backfills of the dorsal neck muscles d1-3 reveal the location of the underlying motor neuronal cell bodies in the suboesophagal ganglion (SEZ), which are shown in close-up (b) (c) In addition to the motor neurons in the SEZ, the neck muscles d1-3 are also innervated by motor neurons in the prothoracic ganglia of the ventral nerve cord (VNC). (d) Dendrites of the neck motor neurons residing in the SEZ extensively arborize in the AMMC region of the brain (e) In the prothoracic ganglion of the VNC, motor neuronal soma send their axons to the muscles through nerves in the prothoracic ganglion. Scale bar in each subpanel is 100 microns.

#### Motor neurons of the ventral and tentorial neck muscles

The ventral muscles v, cv1 and the tentorial muscle t2 in both the left and right halves (inset; Figure 6a) were accessed and filled from the ventral side by removing the ventral sternum cuticle. The motor neurons in the ganglia are bilaterally symmetric and ipsilateral. The neurons in the left and right halves are located both in the SEZ (Figure 6a, b) and prothoracic ganglion (Figure 6a, c), and occur as symmetric pairs. These motor neuronal axons descend through the SEZ and prothoracic ganglionic nerves respectively, to innervate the muscles v and t2. The motor neuronal soma in the SEZ are present in the medial and lateral regions, with dense arborization of dendrites within the AMMC in the deutocerebrum (Figure 6b). The cell bodies are located in the anterior segment of the prothoracic ganglion whereas the highly branched dendritic structures are posterior to the soma (Figure 6c). We also identified ipsilaterally symmetric cell bodies in the ventro-lateral region of SEZ (Figure 6d) as well as prothoracic ganglion (Figure 6e) for cv1 muscle pair.

**Figure 6.**
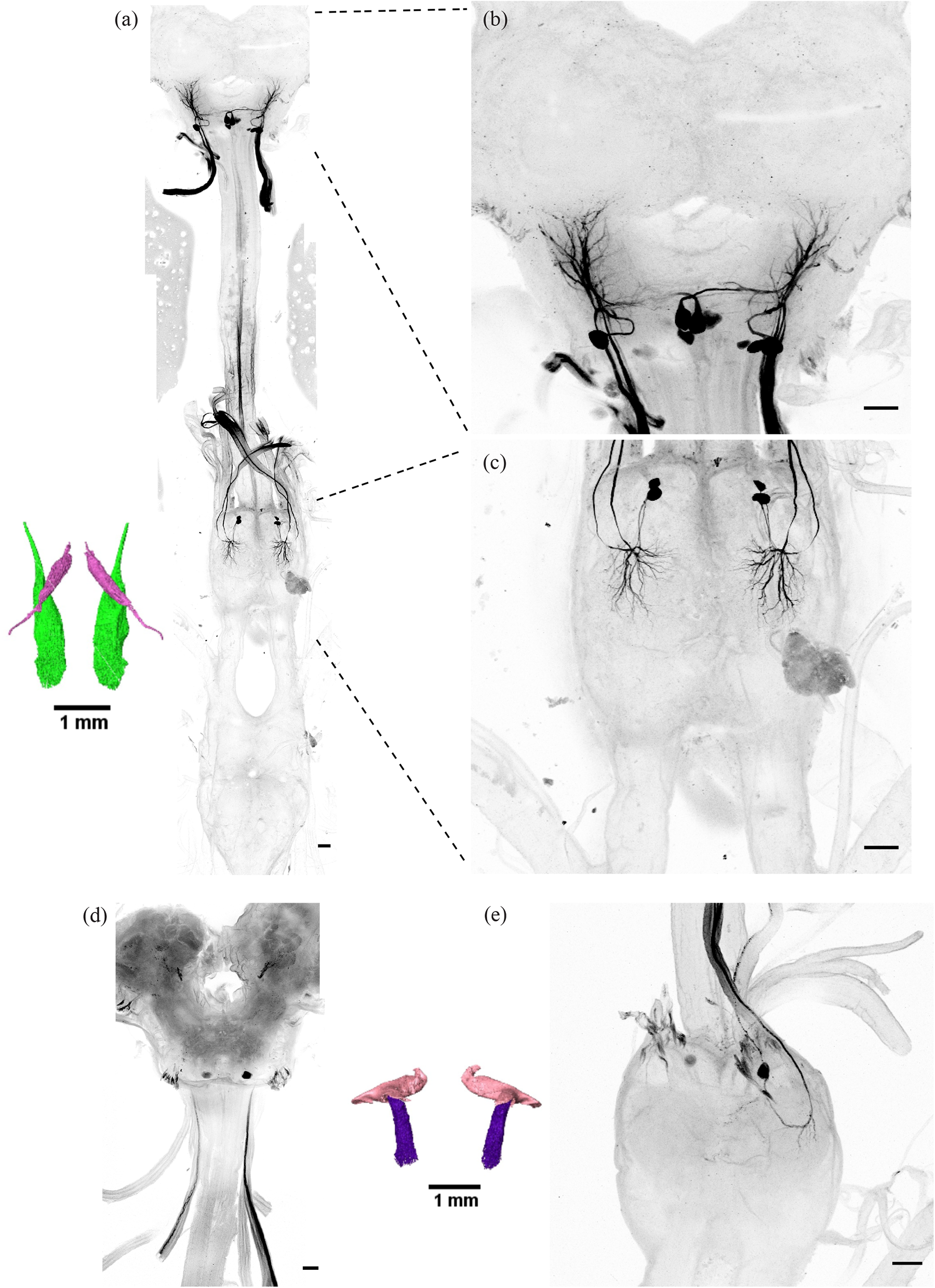
Ventral neck muscle motor neurons. a) Backfills from ventral neck muscle pair v and the ventral tentorial neck muscle pair t2 reveal projections from the neck motor neurons into the central nervous system of moth. Neck muscles articulate into (b) the SEZ region of the brain and (c) the prothoracic ganglion in the ventral nerve chord. (d-e) Backfills from the cv1 neck muscle pair reveal cell bodies of motor neurons in the ventro-lateral region of (d) SEZ and in the anterior part of the (e) prothoracic ganglion. Scale bar in each subpanel is 100 microns.

## 4. Discussion

### 4.1 Neck muscles

#### Action of neck muscles on head movements

The position, attachment points, orientation and geometry of the various muscles outlined in this paper allow us to predict how their activation might actuate head movements. In drawing such predictions however, it is essential to bear in mind that the neck muscle action is combinatorial and muscles function as groups rather than singly (Dickinson *et al* 2000). The hypotheses presented here are thus only first-order hypotheses about the putative function of individual muscles, as summarized in Figure 7 and Table 1. Bilateral action or co-contraction is considered only within single bilaterally symmetric pair of neck muscles.

**Fig. 7.**
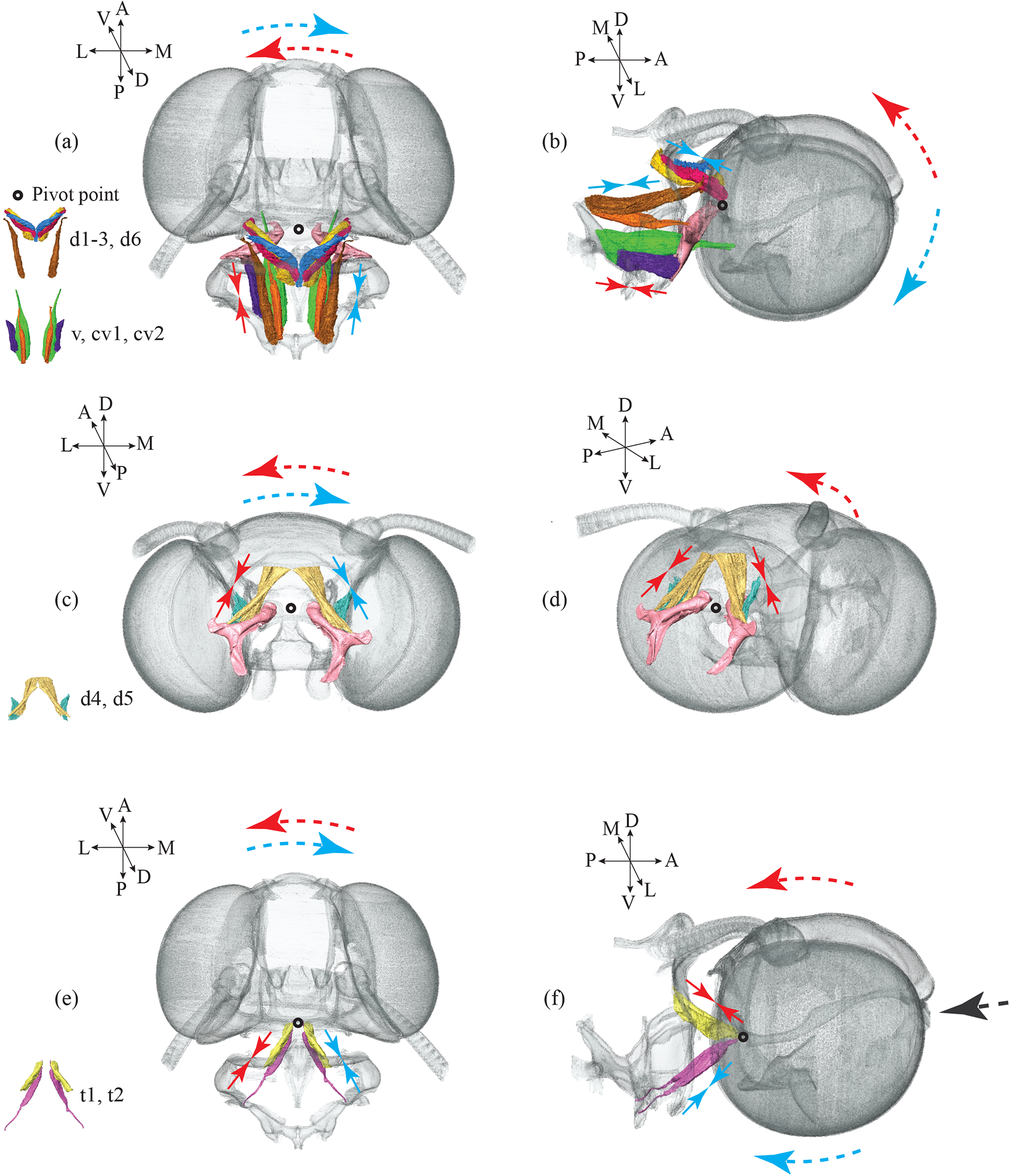
Hypotheses for function of head movements based neck muscle morphology. (a-b) Putative action of the neck muscles d1-3, d6, v, cv1-2 (see inset image for individual muscle identification). (a) Dorsal view of the head with the neck muscles d1-3, d6, v, cv1-2. The contraction of the neck muscles on the left side (red inward-facing arrows) causes counter-clockwise yaw movement of the head (red dashed arrow), whereas contraction of the muscles on the right side (blue arrows) causes a clockwise rotation (blue dashed arrow). (b) Lateral view of the head with the dorso-longitudinal neck muscles shows that co-contraction of the dorsal group of muscles (blue inward-facing arrows) causes a pitch-up (blue dotted arrow), and co-contraction of the ventral group (red inward-facing arrows) causes a pitch-down movement (red dotted arrow) of the head around the pivot point (indicated by a black circle) (c-d) Putative action of the neck muscles (d4-5); see inset image for individual muscle identification). (c) Posterior view of the head with the neck muscles d4 and d5 shows that contraction of the muscles on the left (red inward-facing arrows) causes head to roll counterclockwise (red dotted arrow) whereas on the right (blue inward-facing arrows) causes the head to roll clockwise (blue dotted arrow) (d) Co-contraction of the d4-5 muscles elicits a pitch-up movement of the head (red dotted arrow). This view is slightly tilted from the lateral to provide a clearer image of the bilateral muscles. (e-f) Putative action of the tentorial neck muscles (t1-2) (see inset image for individual muscle identification). (e) Dorsal view of the head with the tentorial muscles t1 and t2 shows that their contraction on the left (red inward facing arrows) causes a counter-clockwise yaw movement of the head (red dotted arrows), whereas contraction on the right (blue inward facing arrows) causes a clockwise yaw movement of the head (blue dotted arrows). (f) Co-contraction of the t1 pair of muscles causes a pitch-up movement (red dotted arrows) and of the t2 group causes a pitch down (blue dotted arrows). If both groups co-contract the head is pulled towards the thorax (black dotted arrow)

**Table 1:** Diverse neck muscles, their organization and predicted head movements.

| Neck muscle pair | Action type | Predicted head movements |
| --- | --- | --- |
| **d1, d2, d3, d6**<br>**(Figure 7a, b)** | On each side | Head yaw |
|  | Co-contraction | Head pitch-up |
| **v, cv1, cv2 (Figure 7a, b)** | On each side | Head yaw |
|  | Co-contraction | Head pitch-down |
| **d4, d5 (Figure 7c, d)** | On each side | Head roll |
|  | Co-contraction | Head pitch-up |
| **t1, t2 (Figure 7e, f)** | On each side | Head yaw |
|  | Co-contraction | Head pulled towards thorax |

| **Neck muscle pair** | **Action type** | **Predicted head movements** |
| --- | --- | --- |
| **d1, d2, d3, d6**<br>**(Figure 7a, b)** | On each side | Head yaw |
|  | Co-contraction | Head pitch-up |
| **v, cv1, cv2 (Figure 7a, b)** | On each side | Head yaw |
|  | Co-contraction | Head pitch-down |
| **d4, d5 (Figure 7c, d)** | On each side | Head roll |
|  | Co-contraction | Head pitch-up |
| **t1, t2 (Figure 7e, f)** | On each side | Head yaw |
|  | Co-contraction | Head pulled towards thorax |

##### Dorsal muscles

From the orientation of dorsal muscles, the following hypotheses can be proposed about their function. First, as the neck muscle pairs d1-3 and d6 are dorso-longitudinally aligned and connect on either side of the neck pivot (Figure 2a-c, g-i), enabling head rotation about the yaw axis. Contraction of muscles on the right would cause a clockwise yaw rotation of the head (blue arrows, Fig 7a) whereas contraction on the left would cause an anticlockwise rotation (red arrows, Fig 7a). Moreover, because these muscles attach above the pivot point, their co-contraction would cause the neck to pitch up (red arrows, Figure 7b). Second, the dorso-ventral neck muscle pairs d4-5 are anchored laterally outward about the pivot point on the cervicale (Figure 2d-f), and would cause the head to perform roll movements (Figure 7c). Thus, contraction of these muscles on the left side (red arrows) would cause counter-clockwise, and on the right side (blue arrows) would cause clockwise rotations of the head (Figure 7c). Third, as these muscles run between the cervicale and the post-occipital ridge of the head, their co-contraction would thus cause the head to pitch up (black arrows, Figure 7d).

##### Ventral muscles

The ventral muscles v, cv1-2 (Figure 3; Fig 7a-b) have previously been hypothesized to actuate yaw head rotations in the silkmoth (Mishima & Kanzaki, 1998). In the Oleander hawk moth, the ventral muscle v (Figure 3a) is the thickest neck muscle with sharply tapering slope and conical geometry which focuses its force on the ventrolateral end of foramen magnum. Their action on each side will enable strong head-movements around the yaw axis (Figure 7a). Because the muscle pairs are laterally outward and below the pivot point, their co-contraction would cause the head to pitch-down (blue arrows, Figure 7b). Because the ventral neck muscles cv1-2 (Figure 3d-f) which pull the head *via* the cervicale also have similar orientation as the ventral muscle v, they too will contribute to the yaw motion (Figure 7a) upon alternating action and pitch-down rotations of the head upon co-contraction (Figure 7b).

##### Tentorial muscles

The tentorial muscles t1-2 (Figure 7e, f) attach medially and centrally at the tentorial bridge which is also the pivot-point. Their contraction alternatingly on the left (red arrows) and right (blue arrows) would cause the head to yaw with respect to thorax (Figure 7e). However, their co-contraction would pull the head closer to the thorax (Figure 7f). These hypotheses are summarized in Table 1.

#### Direct and indirect neck muscles

Functionally, the neck muscles may be classified as either *direct* and *indirect*. The direct muscles include the dorso-longitudinally oriented dorsal muscles (d1-3; Figure 2 a-c), the ventral muscle v and the tentorial neck muscles t1, t2 (Figure 4). These exert forces directly on the head while being anchored on sclerites in the prothoracic skeleton. On the other hand, the indirect neck muscles exert forces on the head *via* the cervical sclerite. We identified two subtypes – the internal-indirect (d4, d5; Figure 2 d-f) which are anchored between the head and the cervicale whereas the external-indirect neck muscles (cv1, cv2; Figure 3 d-f) span between the cervicale and the prothoracic cuticle (nomenclature adapted from Berry & Ibbotson, 2010). The internal-indirect muscles d4, d5 have their base anchored on the anterior part of the cervicale but the external-indirect muscles cv1, cv2 attach to the posterior part of the cervicale and exert forces from behind. Thus, a subset of the neck muscles can directly rotate the head whereas the other subset creates indirect action on the head-movements via the cervicale.

### 4.2 A comparative view of the neck skeletal structures

The structural architecture of the insect neck skeleton has been previously described in a few insect orders including flies (Strausfeld, Seyan & Milde, 1987), bees (Berry & Ibbotson, 2010), locusts (Shepheard, 1974) and moths (Eaton, 1971). In all cases, the outer covering of the neck is soft and flexible, which enables free movement of the head relative to thorax. Across insects, the bilaterally symmetric cervical sclerite or cervicale, is shaped like a pair of lever arms with varying shape and size. In the hawkmoth *Daphnis nerii*, the cervicale provides the attachment point for four pairs of neck muscles and is thus a key component of the neck movement apparatus. Its structure has features that are interesting from a mechanical perspective. For instance, it appears to have a decreasing gradient of sclerotization from the prothoracic end towards the posterior head capsule both in moths (Supplementary Figure 2) and blowflies (Strausfeld, Seyan & Milde, 1987). Further research is needed to examine the biomechanical implications of this gradient of sclerotization.

The force generated by contraction of the neck muscles acts either directly on the posterior capsule of head cuticle, or indirectly *via* the cervicale. This should cause the head to rotate about the anterior tip of the cervical sclerites which acts as a pivot. The occipital condyle process – the socket into which the cervicale inserts – serves as a partial rigid pivot point for rotation of the head capsule (Figure 1b, c). The condyle action has been described in flies (Sandeman & Markl, 1979) and the direct or indirect neck muscles action in bees (Berry & Ibbotson, 2010), but it is not clear how general these structures are across insects. In a horizontal two-point pivot system, the head can readily undergo pitch movements, but not roll or yaw movements which involve tilting the tentorial bridge along the dorso-ventral axis. For yaw movements of the head, one of the occipital condyle processes must move either forward or backward relative to the other. Similarly, for roll movements, one of the occipital condyle processes must go up or down relative to other, about the horizontal tentorial axis. These might be possible if the cervicale is flexible with respect to the occipital condyles in the head as it is least sclerotized in the anterior tip, but rigid at the prothoracic attachment sites due to heavy sclerotization (Supplementary Figure 2c, d). In insects including blowflies and crickets, the tentorial bridge is very small, and the two pivot points for the cervicale arm pairs act as if they are fused as one pivot point. The engagement and disengagement dynamics between the anterior tip of the cervicale arms and their sockets may also be interesting to investigate. The mechanics of head movements thus arise from a combinatorial activity in the neck muscles acting *via* the rigid cervicale. To understand head movements, it is thus important to consider the internal structure of the neck joint at the posterior section of the head capsule, its connection to the prothorax (sternal cuticle) for stability, and the related mechanics (Snodgrass, 1928, 1960, 2018).

### 4.3 Neck motor neurons

Like other invertebrate muscles, a neck muscle in *Daphnis nerii* may receive innervation from multiple motor neurons, but each motor neuron specifically innervates only one muscle. Because the neck muscles span the head-neck joint, the action of each muscle must be coordinated by motor neurons that reside either in the sub-esophageal zone (SEZ), or in the pro-thoracic ganglion, or both, with their axonal tracts lying in multiple efferent nerves emanating from the SEZ and the prothoracic ganglia. The dorsal neck muscles (d1-3) which are putatively involved in yaw or pitch up motions of the head (Figure 7a, b) are innervated by multiple motor neurons from two ganglia *viz* SEZ and prothoracic ganglion (Figure 5a-e). The ventral muscle cv1 which may be involved in yaw or pitch down motions of the neck (Figure 7a, b) is innervated by only a single ipsilateral motor neuron in the SEZ (Figure 6c).

However, the muscles (v) which are putatively involved in yaw, pitch-down (Figure 7a, b) and t2 which may help pull the head towards thorax (Figure 7f), are innervated by neurons residing in the SEZ and the prothoracic ganglion (Figure 6a-c). Almost all neck muscles are innervated by ipsilateral motor neurons, with the sole exception of the muscle t2 which is innervated by a contralateral motor neuron cell body located medially in the SEZ below the foramen but their dendrites and axon terminals are ipsilateral with the muscle (Figure 6b).

Notwithstanding the differences in the overall neck muscle morphology of diverse insects, there are some common themes in their motor neuronal architecture. In locusts, the neck region contains 14 head-neck muscle pairs, innervated by approximately 60 axons of motor neurons from the SEZ, prothoracic and mesothoracic ganglia (Shepheard, 1973, 1974). The muscle fiber types are characterized based on their excitability, decay time constants in the spikes and whether the type of axon innervating the muscle was tonic (usually fast), phasotonic (usually intermediate) or phasic (usually slow) (Shepheard, 1973). A comparison of the neck motor system between locusts and crickets (both Orthoptera) show that the location of motor neurons in ganglia and the morphology of the neck motor neurons are similar (Honneger *et.al.* 1984, Altman & Kien, 1979). In honey bees (Hymenoptera), there are 17 pairs of neck muscles, along with their attachments and the relevant skeletal structures connecting posterior head capsule and prothorax for head-neck joint articulation *via* the cervical structures (Berry & Ibbotson, 2010). The blowfly neck consists of 21 bilaterally symmetric muscle pairs and here too the cervicale acts as the anchoring structure for the arthrology of head-neck joint articulation (Strausfeld, Seyan & Milde, 1987; Milde, Seyan & Strausfeld, 1987).

The general organization of neck motor neurons in the moth brain is similar to bees (Schröter *et.al*., 2007; Goodman, Fletcher *et.al*., 1987; Hung, Kleef & Ibbotson, 2011) and flies (Strausfeld, Seyan & Milde, 1987; Milde, Seyan & Strausfeld, 1987; Kauer, Borst & Haag, 2015). The dendritic arborizations are dense and elaborate in the deutocerebrum and tritocerebrum spanning the AMMC with the cell bodies located either medially, below the foramen or laterally in the SEZ (Figure 5b, d, Figure 6b). Motor neurons in prothoracic segment of the nervous system have cell bodies located anterior and medial in the ganglion whereas their dendritic ramifications are posterior to the soma (Figure 5e, 6c, e). These locations in the SEZ or prothoracic ganglion receive a wide variety of sensory inputs, including mechanosensory inputs from the Johnston’s organs (Dieudonné, Daniel & Sane, 2014, Sant & Sane, 2018) and cephalic hair (Manjunath & Sane, *pers. comm*) to which they may be connected either directly or indirectly. This possibility needs to be further explored using double fills of sensory and motor neurons, and electrophysiological recordings from neck muscles.

### 4.4 Multisensory integration and the neck motor system

To generate head movements in diverse behavioral contexts, the neck motor system in insects integrates multiple sensory inputs (Strausfeld & Seyan, 1985). These include optic flow from the compound eyes (moths, Dombrowski, Milde & Wendler, 1990; flies; Gronenberg, Milde & Strausfeld, 1995; Wertz, Haag & Borst, 2012), olfactory descending neurons in moths (Kanzaki & Mishima, 1996), mechanosensory feedback from halteres in Diptera (Sandeman & Markl, 1979; Tracey, 1975; Huston & Krapp, 2009), antennae in moths (Chatterjee et al, 2022), cerci, antennae and legs in crickets (Horn & Bischof, 1982) and wings in moths (Ando et.al. 2011). In blowflies, the haltere feedback modulates neck motor neurons at latencies of 2.5-3 ms (Sandeman & Markl, 1979), on the order of single wing strokes. In addition, the neck motor system also responds strongly to inputs from the mechanosensory prosternal organs (Preuss & Hengstenberg, 1992; Hengstenberg, 1993). In Diptera, the prosternal organ (Preuss & Hengstenberg, 1992; Paulk & Gilbert, 2006) consists of two sets of symmetric hair fields that are in apposition to one another at the base of the neck, and senses instantaneous head position relative to thorax. These sensory afferents terminate in the prothoracic ganglion which also contain the dendrites of the neck motor neurons in close proximity suggesting rapid interaction between sensing head orientation and actuating head movements. The neck prosternal chordotonal organ (pCO) is composed of several sensory axons that end in the median ventral association center of various neuromeres within the thorax-abdominal ganglion (Stölting, Stumpner & Lakes-Harlan, 2007) of the sarcophagid fly *Sarcophaga bullata*. These cells contain receptor cells with dorso-lateral branches in the mesothoracic neuromere which are insensitive to frequencies below approximately 1 kHz, and receptor cells without such branches, respond to lower frequencies (Stölting, Stumpner & Lakes-Harlan, 2007). They are thought to encode head rotations and foreleg vibrations which are transduced to the attachment site of the pCO (Stölting, Stumpner & Lakes-Harlan, 2007).

In addition to inputs from the prosternal organ, the axons of descending motion-sensitive visual interneurons (Gilbert et al 1995; Gronenberg et al 1995; Goodman *et.al*., 1987; Milde et al 1987; Gronenberg & Strausfeld, 1991; Gronenberg & Strausfeld, 1990; Milde & Strausfeld, 1986; Wertz et al 2012; Strausfeld & Bassemir, 1985; Kauer et al 2015) are surrounded by the dendrites of neck motor neurons in both brain and prothoracic ganglion, suggesting direct synapses. In honey bees, the axonal projections of hair receptor fields perhaps homologous to the pro-sternal organ, likely encodes head positioning relative to body (Ai & Hagio, 2013) and ascends from the thoracic ganglia to the AMMC and SEZ. In contrast,the descending antennal mechanosensory afferents arborize in the AMMC, close to the dendrites of neck motor neurons. The prosternal organ has not so far been reported in Lepidopteran insects, and detailed anatomical studies are required to identify its location.

Together, these papers suggest that optic-flow cues from the visual system (eyes), air-flow and tactile cues from mechanosensors (cephalic and cerci hairs, antennal Johnston’s organs), vestibular cues from mechanosensors (haltere and wing campaniforms, antennal Johnston’s organs) and proprioceptive feedback from mechanosensors (prosternal organ, pCO, neck hair plates) interact with the neck motor apparatus in insects.

### 4.5 Neck motor system and gaze stabilization

Neck movements are essential in all forms of insect locomotion due to their central role in gaze stabilization (e.g. Land, 1999). To generate a stable representation of the external world on their retina, flying insects rely on optomotor reflexes (Borst, 2014; Haag, Wertz & Borst, 2010; Goodman, 1964; Collett & Land, 1974, 1975; Kern & Varju, 1998) which includes compensatory head movements (Hengstenberg, Sandeman & Hengstenberg, 1988). Through the combinatorial action of multiple neck muscles, insects perform smooth compensatory head movements to ensure minimization of optic flow (i.e. gaze stabilization) during body rotations (Hardcastle & Krapp, 2016). Several neck motor neurons located in the SEZ respond to wide-field visual stimuli over compound eyes and receive inputs from motion-sensitive descending visual interneurons (Hung, Kleef & Ibbotson, 2011; Schröter *et.al*., 2007). Because flight maneuvers are rapid, visual feedback alone may be insufficient as guiding cues for the compensatory head movements. Similar to the vestibulo-ocular reflexes in vertebrates which integrate both visual feedback from eyes and vestibular cues from the inner ear system (e.g. Fetter, 2007), insects also rely on visual cues from their compound eyes and vestibular feedback from either their halteres (reduced hind wings of Diptera; Pringle, 1948) or from the antennal mechanosensory Johnston’s organs (in Lepidoptera; Sane *et al*, 2007; Dahake *et al*, 2018; Chatterjee *et al*, 2022; also see Sane et al 2023). Thus, to generate compensatory head movements for stabilizing gaze, the neck motor system integrates information primarily from visual and mechanosensory modalities (Hengstenberg, 1993; Chatterjee *et.al*. 2022). Moreover, recent studies have also showed that when head-movements are restricted in moths, they are unable to control their flight (Chatterjee *et.al*., 2022). In Diptera, restricted head movements in tethered Drosophila caused reduced thrust forces for lift (Cellini & Mongeau, 2020). Moreover, integration of visual and haltere mechanosensory cues is non-linear in flies (Huston & Krapp, 2009) but, in moths and many other insects, the mechanistic details of how the antennal mechanosensory and visual feedback are integrated at the neck motor is an open question. The data presented in this paper will be useful for further studies on the role of neck muscles in gaze and flight stabilization.

## Acknowledgments

We acknowledge the Micro-CT imaging facility at NCBS, Bengaluru, India and Mr. Sunil Prabhakar for help with the X-ray scanning microscopy, Abin Ghosh for help with Amira software, Girish Kumar G. S. for help in making the interactive Adobe pdf. We also acknowledge Maitri M. for help with neuroanatomy and the Central Imaging and Flow Cyctometry Facility (CIFF) at NCBS for the confocal microscopy imaging. Dr. Harshada H. Sant helped with comments and discussions on the manuscript. Funding was provided by the Air Force Office of Scientific Research grants (AFOSR; FA2386-11-1-4057 and FA9550-16-1-0155), Human Frontiers Science Program (RGP0066/2012) and the Tata Institute of Fundamental Research, National Centre for Biological Sciences, Department of Atomic Energy, Government of India to SPS.

## Supplementary figures

**Supplementary Figure 1. Interactive 3D model**

An interactive 3D model of the neck musculo-skeletal apparatus in the Oleander hawkmoth which can be accessed through Adobe pdf reader. Various skeletal structures and muscles can be selectively displayed using the “Toggle Model tree” option.

**Supplementary Figure 2. Neck cervicale**

(a) Occipital condyle as the sockets for cervicale in the posterior head capsule near the tentorial bridge.

(b) Dorsal view of the cervicale pair anchored in their sockets. The attachment of the anterior tip of the cervicale at these sockets act as the pivot point of the head with respect to neck.

(c) Dorsal view of the entire thorax with cervicale pair protruding out of the prothorax and their anterior tip exposed as the head capsule is removed.

(d) Decreasing gradient of sclerotization towards the anterior tip of the cervicale where it attaches the head.

(e) Oblique dorsal side view showing the attachment of the cervicale pair onto the head capsule.

(f-h) Whitish coloration in the anterior tip of the cervicale (within pink square) indicate change in cuticular composition and fluorescence with ultraviolet excitation may indicate the presence of resilin or resilin-like protein

**Supplementary Figure 3. Neck sternum**

(a) Lateral side view of the prothoracic skeleton showing (b) sternum in green and the Furca I in pale yellow

(c) Posterior view of the prothoracic skeleton showing (d) sternum in green and the Furca I in pale yellow

(e) Ventral and (f) dorsal view of the sternum cuticle

## Supplementary Materials

### Recipe for insect saline

The base solvent for the salts is distilled water.

Stock solution – 1 is made using 3.8gm/L NaCl, 24.97g/L KCl and 32.7333g/L MgCl_2_. 6H_2_O

Stock solution – 2 is made with 1.8g/L Na_2_HPO_4_

Stock solution – 3 is made using 1.95g/L NaH_2_PO_4_

Stock solution – 4 is made with 6.175g/L CaCl_2_ .2H_2_O

For making 500mL of 1x insect saline:

1. Add 50 ml of stock solution 1; 50ml of stock solution 2 and 50ml of stock solution 3. Mix it well

2. Add 50 ml of stock solution 4 to the above solution dropwise slowly with continuous mixing

3. Adjust the pH of the solution to 6.7

4. Add 172.9 mM of Dextrose – 15.57g/500 ml

5. Make up to 500 mL using distilled water

#### Recipe for Phosphate Buffer Solution (PBS)

To make 20 ml of 10x PBS or 200 ml of 1x PBS, add 0.0512 g of NaH _2_ PO _4_, 0.238 g of Na _2_ HPO _4_ and 2.044 g of NaCl in 200 ml of distilled water. Adjust the pH of the solution to 7.4

#### Recipe for Paraformaldehyde (PFA)

To make 100 ml of 4% paraformaldehyde fixative solution, take 4 g of PFA and add it to a container with 100mL of 1x PBS. Stir this mixture on a magnetic stirrer at 65-70°C for 3.5 hours in the fume hood. For best results, use the PBS based PFA solution within 14 days for neuroanatomy.

## Notes

### Competing Interest Statement

The authors have declared no competing interest.

