## Supplementary figures and images for "The motor apparatus of head movements in the Oleander hawkmoth (*Daphnis nerii*, Lepidoptera)"

### Supplementary Figure 1

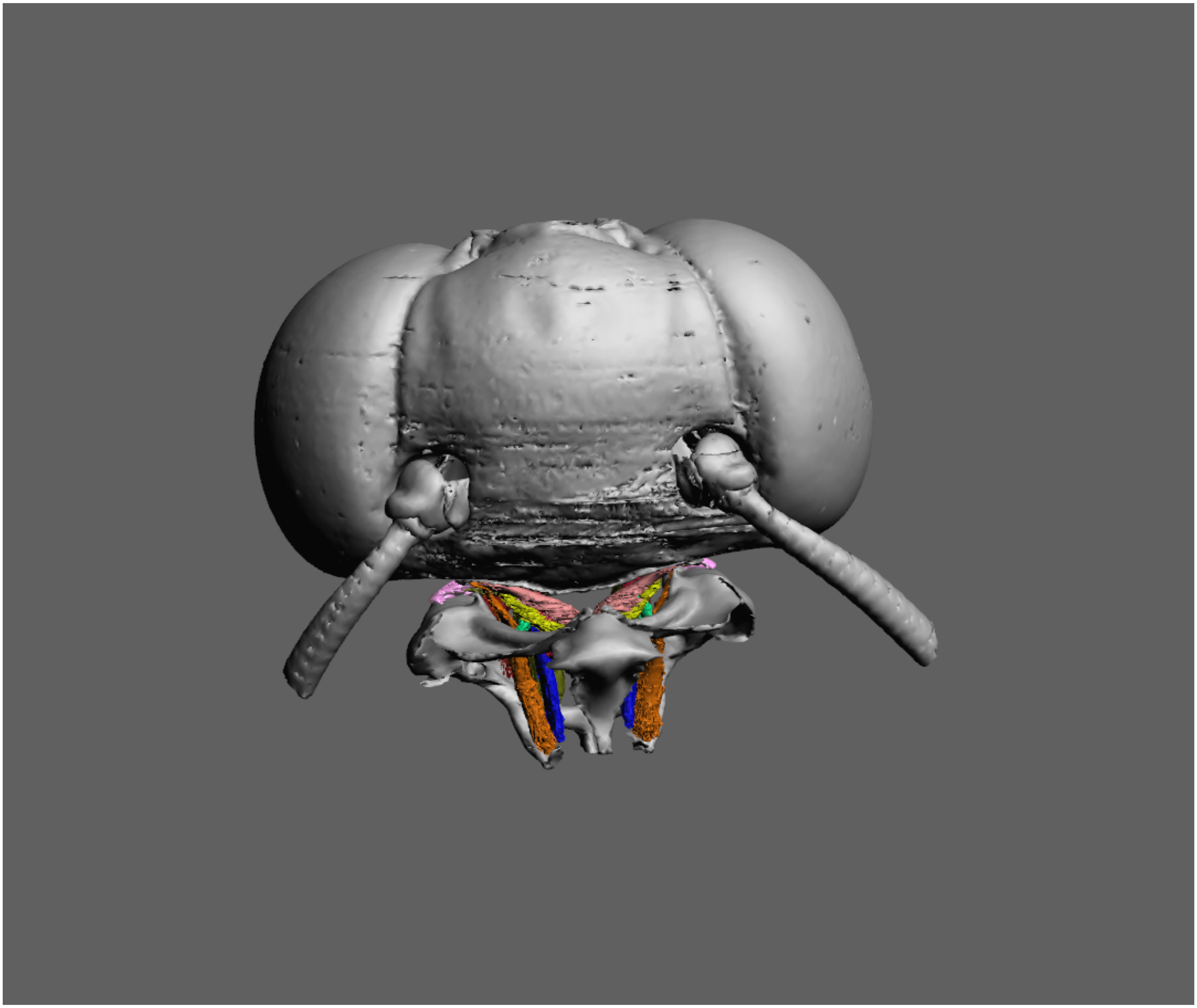

### Supplementary Figure 2

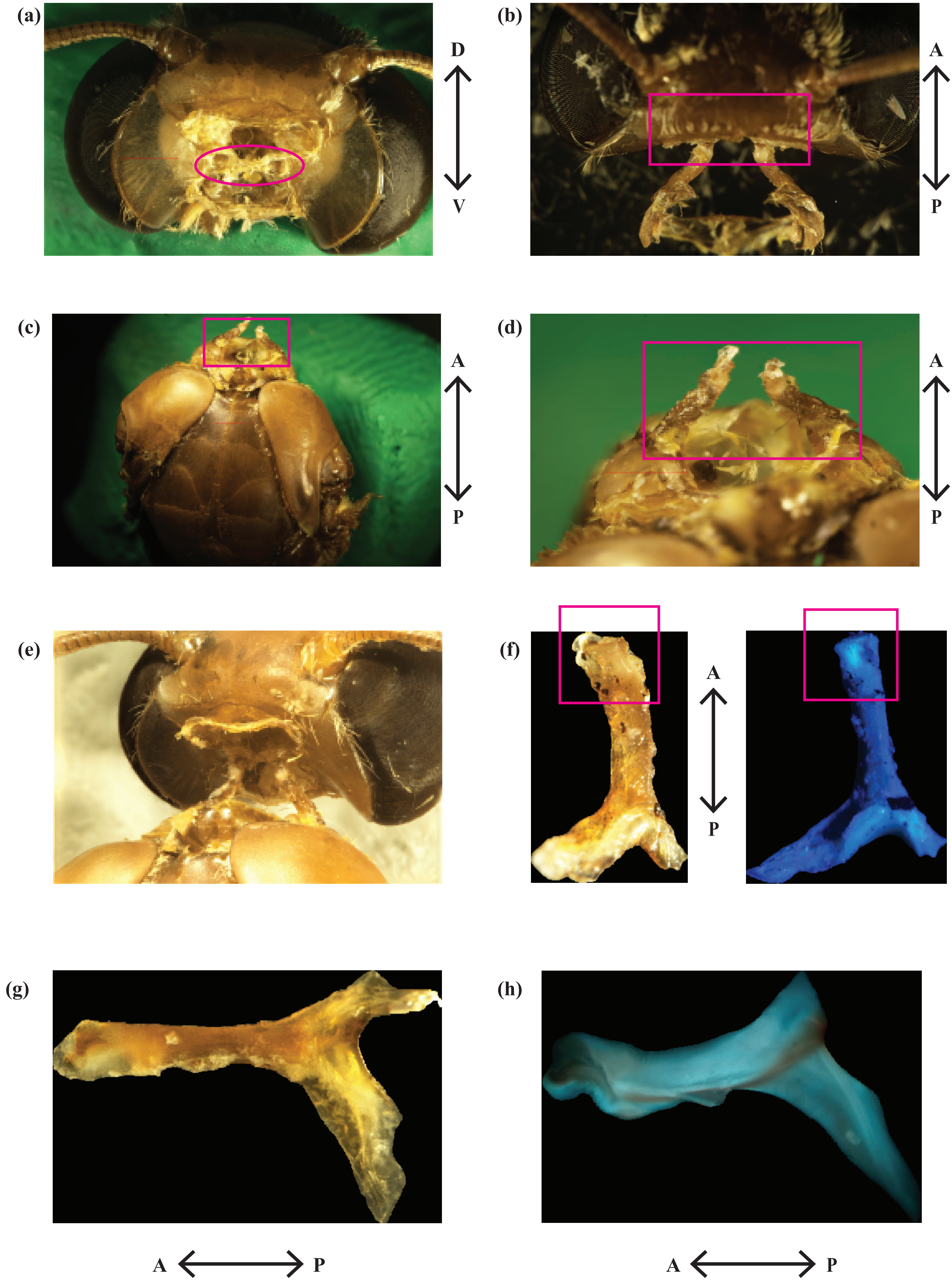


**Supplementary Fig. 2. Neck cervicale**

### Supplementary Figure 3

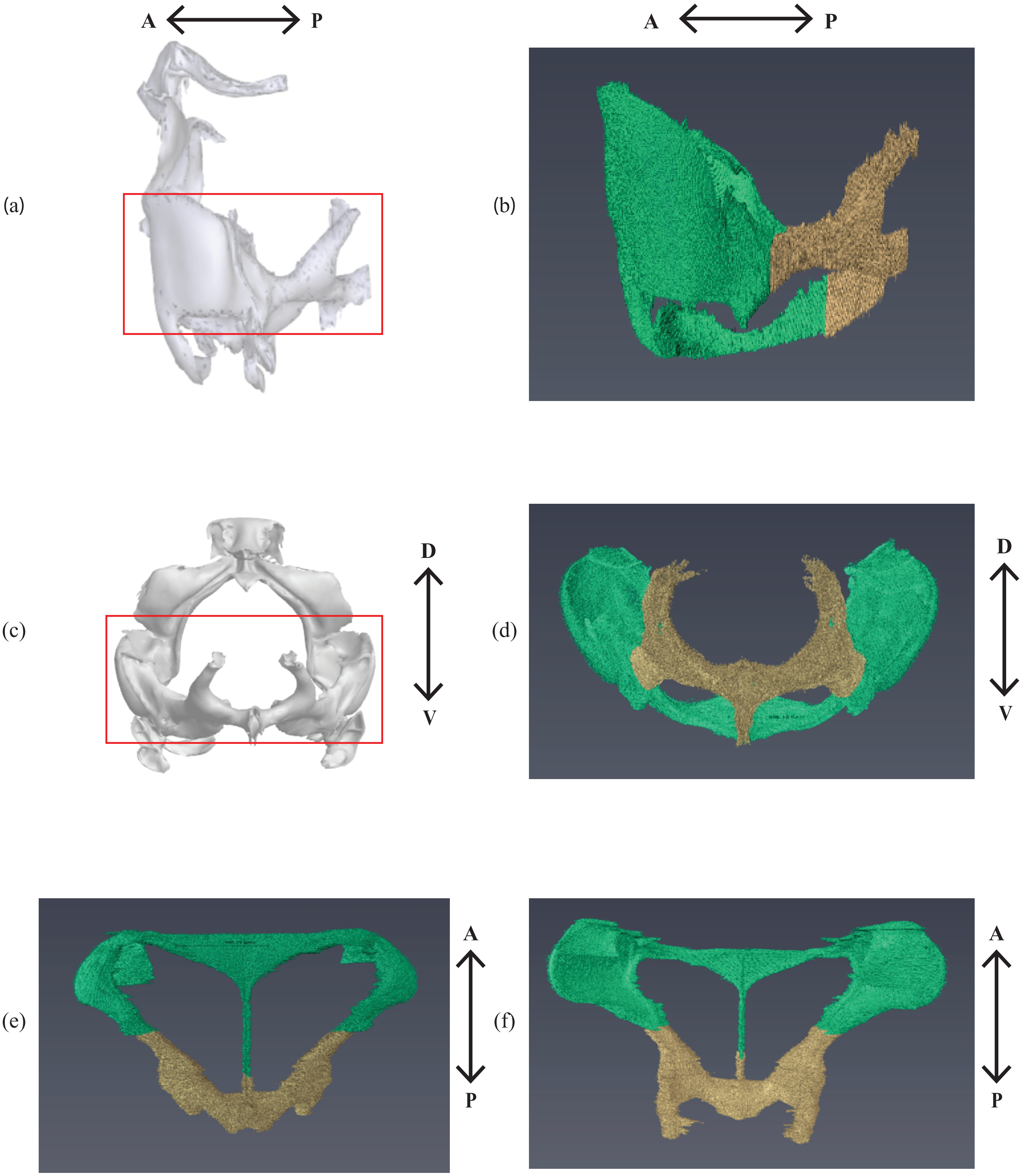


**Supplementary Fig. 3. Neck sternum**
